# Cortical processing of chemosensory and hedonic features of taste in active licking mice

**DOI:** 10.1101/2020.02.11.923565

**Authors:** Cecilia Bouaichi, Roberto Vincis

**Affiliations:** Department of Biological Science, Florida State University, Tallahassee, FL, 32306, USA.; Program in Neuroscience, Florida State University, Tallahassee, FL, 32306, USA.

**Keywords:** Extracellular recording, GC, gustation, active licking, coding, taste.

## Abstract

In the last two decades, a considerable amount of work has been devoted to investigating the neural processing and dynamics of the primary taste cortex of rats. Surprisingly, much less information is available on cortical taste electrophysiology in awake mice, an animal model that is taking a more prominent role in taste research. Here we present electrophysiological evidence demonstrating how the gustatory cortex (GC) encodes information pertaining the basic taste qualities (sweet, salty, sour, and bitter) when stimuli are actively sampled through licking, the stereotyped behavior by which mice control the access of fluids in the mouth. Mice were trained to receive each stimulus on a fixed ratio schedule in which they had to lick a dry spout six times to receive a tastant on the seventh lick. Electrophysiological recordings confirmed that GC neurons encode both chemosensory and hedonic aspects of actively sampled tastants. In addition, our data revealed two other main findings; GC neurons encoded information about taste identity in as little as 120 ms. Consistent with the ability of GC neurons to rapidly encode taste information, nearly half of the recorded neurons exhibited spiking activity that was entrained to licking at rates up to 8 Hz. Overall, our results highlight how the GC of mice processes tastants when they are actively sensed through licking, reaffirming and expanding our knowledge on cortical taste processing.

**NEW & NOTEWORTHY:** Relatively little information is available on the neural dynamics of taste processing in the mouse gustatory cortex (GC). In this study we investigate how the GC encodes information of the qualities and hedonics of a broad panel of gustatory stimuli when tastants are actively sampled through licking. Our results show that the GC neurons broadly encode taste qualities but also process taste hedonics and licking information in a temporally dynamic manner.

## INTRODUCTION

The gustatory cortex (GC) is the primary cortical region responsible for processing taste information (Spector and Travers 2005; Carleton et al. 2010; Vincis and Fontanini 2019). Over the past decades, many studies have investigated the neural representation of gustatory stimuli in the GC of alert rats, an animal model that has been extensively used for the psychophysical examination of taste (Spector and Travers 2005; Blonde et al. 2015). Electrophysiological analysis of spiking activity has revealed that the GC of rats encodes multiple facets of taste experience (Carleton et al. 2010; Maffei et al. 2012; Vincis and Fontanini 2016), including the physiochemical (Katz et al. 2001; Stapleton et al. 2006) and psychological (Katz et al. 2001; Grossman et al. 2008; Jezzini et al. 2013; Mukherjee et al. 2019) aspects of gustatory stimuli, as well as the expectation of taste (Stapleton et al. 2006; Saddoris et al. 2009; Samuelsen et al. 2012).

Relatively less work has been done in mice, an animal model that offers easier access to genetic tools to manipulate and visualize neuronal activity. Although a considerable amount of studies have investigated either spatial features of cortical taste-evoked activity *in-vivo* (Chen et al. 2011; Fletcher et al. 2017; Lavi et al. 2018; Livneh et al. 2017) or taste behavior (Graham et al. 2014; Peng et al. 2015), limited information is available on cortical taste electrophysiology in awake mice. One recent study described how GC neurons encoded gustatory information when taste stimuli were injected into the mouths of alert mice via intra-oral cannulas (IOCs) (Levitan et al. 2019). While IOCs provide a reliable and rapid method to deliver taste solutions, they add a degree of passivity to taste delivery and could potentially alter the sequence of events associated with the neural processing of gustatory information (Roussin et al. 2012). In contrast, liquid gustatory stimuli are normally sensed by rodents through licking - a stereotyped behavior by which fluids are actively introduced in the mouth (Travers et al. 1997; Graham et al. 2014). However, several key issues regarding cortical taste processing in active licking mice are largely unaddressed. Specifically, in this study we investigated two important questions.

The first question reverts around the temporal evolution of taste-evoked neural representations. Previous experiments using IOCs showed that taste processing within the GC is characterized by a dynamic and time-varying modulation of the firing activity extending up to multiple seconds after stimulus delivery (Levitan et al. 2019). However, in active licking mice and rats, smaller post-taste intervals (up to 0.5 s) are normally used to evaluate the processing of chemosensory coding (Stapleton et al. 2006; Vincis et al. 2019). As a result, it is not known if and how gustatory information of the four basic taste qualities evolves over a longer time interval after taste detection by active licking mice.

The second question pertains to the processing of hedonic value (i.e. if a taste is palatable or unpalatable). Previous data have indicated that GC neurons not only encode the chemosensory identity of tastants but also code for palatability, a feature normally emerging 0.5 s after stimulus detection (Katz et al. 2001; Mukherjee et al. 2019; Levitan et al. 2019). While these studies used gustatory stimuli delivered via IOCs, it still unknown whether GC neurons encode hedonic value of tastants even when actively sampled via licking.

To address these questions, we designed and conducted experiments to determine the electrophysiological features of cortical processing of the four basic taste qualities (sweet, salty, bitter, and sour) and water associated with active sensing in mice across a long post-stimulus temporal window. Spiking activity was recorded from single GC neurons in mice permitted to freely lick to receive tastants. To separate neural activity evoked by the gustatory stimuli from electrophysiological correlates of sensory and motor aspects of licking, we 1) trained the mice to receive each taste on a fixed ratio schedule and 2) did not start recording neural activity until the licking pattern evoked by each gustatory stimulus was similar across a 1.5 s temporal window. As a result, the neural response evoked by the tastants could be compared with the one elicited by licking the dry spout before taste delivery. This also served to make sure that neural activity evoked by the distinct tastants would not be impacted by differences in taste-evoked licking variables such as inter-lick interval and lick numbers.

Overall, our results are consistent with previous experiments involving IOCs in both mice and rats, indicating that GC neurons recorded from active licking mice encode both the chemosensory and hedonic aspects of gustatory stimuli in a broad and temporally rich fashion. Additionally, our data provide two additional insights into GC processing in mice. First, the neural activity of cortical neurons encoding taste information can be also modulated by and be coherent with licking. Second, the temporal dynamics of taste response are fast; different from reports using IOCs, taste identity pertaining all taste qualities is coded within 320 ms, and palatability information arises within 1 s post taste delivery. In summary, our data obtained from mice actively sensing a broad panel of gustatory stimuli, confirm and significantly expand our knowledge on cortical taste processing.

## MATERIAL AND METHODS

### Experimental subjects

The experiments in this study were performed on 4 male and 2 female wild type C57BL/6J adult mice (10-20 weeks old). Mice were purchased from The Jackson Laboratory (Bar Harbor, ME); upon arrival, mice were housed on 12h/12h light-dark cycle and had *ad libitum* access to food and water. Experiments and training were performed during the light portion of the cycle. Six days before training began, mice were water restricted and maintained at 85% of their pre-surgical weight. All experiments were reviewed and approved by the Florida State University Institutional Animal Care and Use Committee (IACUC) under protocol #1824.

### Surgery and tetrode implantation

Prior to surgery, mice were anesthetized with a mixture of ketamine/dexmedetomidine (13.3 mg/ml; 0.16 mg/ml). The depth of anesthesia was monitored regularly via inspection of breathing rate, whisking, and periodically estimating the tail reflex. Additional ketamine was supplemented by 30% of the induction dose as needed throughout the surgery. A heating pad (World Precision Instruments) was used to maintain body temperature at 35°C. After the achievement of surgical level of anesthesia, the animal’s head was shaved, cleaned and disinfected (with iodine solution and 70% alcohol), and fixed on a stereotaxic holder. A first craniotomy was drilled above the left GC on the mouse’s skull (AP: 1.2 mm, ML: 3.5 mm relative to bregma) to implant a movable bundle of 8 tetrodes and one single reference wire (Sandvik-Kanthal, PX000004) with a final impedance of 200-300 kΩ for tetrodes and 20-30 kΩ for the reference wire. A second hole was drilled on top of the visual cortex where a ground wire (A-M system, Cat. No. 781000) was lowered ∼300*µ*m below the brain surface. During surgery, the tetrodes and reference wires were lowered 1.2 mm below the cortical surface; they were further lowered ∼200 *µ*m before first day of recording and ∼80 *µ*m after each recording session. Before implantation, tetrodes wires were coated with a lipophilic fluorescent dye (DiI; Sigma-Aldrich), allowing us to visualize the final location of the tetrodes at the end of each experiment. Tetrodes, ground wires, and a head screw (for the purpose of head restraint) were cemented to the skull with dental acrylic. Animals were allowed to recover for a week before the water restriction regimen began. For 3 consecutive post-surgery days, we administered a subcutaneous injection of carprofen (5 mg/kg) to reduce pain and inflammation.

### Taste delivery system and licking detection

The taste delivery system consisted of five separate taste lines (four for tastants and one for water) that converged at the tip of the licking spout. The licking spout contained independent polyimide tubes (MicroLumen, ID = 0.0142), each one connected to one taste line. Gustatory stimuli (sucrose [200 mM], NaCl [50 mM], citric acid [10 mM], quinine [0.5 mM]; from Sigma-Aldrich further dissolved in water to reach the final concentration and presented at room temperature) and water were delivered via gravity by computer controlled (Bpod, Sanworks) 12 V solenoid-valves (LHDA1231115H, Lee Company) calibrated to deliver a 3 *µ*l droplet of fluid (in the context of our rig, the opening times of the solenoid valves to deliver 3 *µ*l of fluid ranged between 15 and 24 ms). Each lick was detected when the tongue crossed an infrared light beam (940 nm) positioned just in front of the drinking spout (Fig. 2A). The beam was generated by a fiber-coupled LED (M940F1, Thorlabs) and received by a photodiode (SM05PD1A, Thorlabs). Lick and taste delivery timestamps were recorded and synchronized with neural data acquisition (Plexon system, see below for more details) and Bpod.

**Figure 1.**
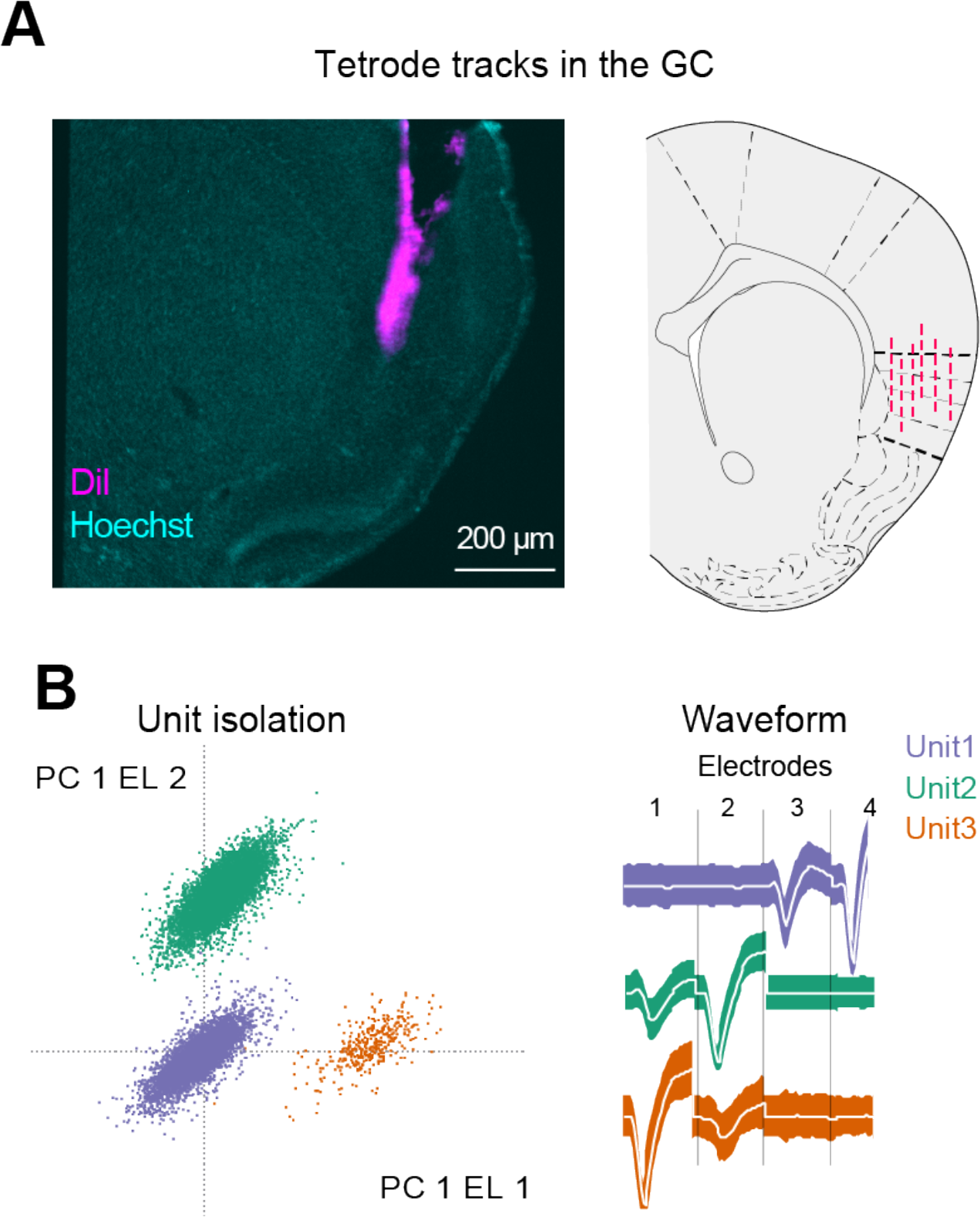
Tetrodes placement and single neuron recording. *A: Left,* example of histological section showing the tetrode tracks (magenta) in the left GC; *right,* schematic of the summary of the tetrodes tracks from the six mice. *B: Left,* representative single unit recordings in the GC showing the principal component analysis (PCA) of waveform shapes for spikes of three individual neurons; *right*, average single unit spike responses for the same three neurons, recorded from the four electrodes.

**Figure 2.**
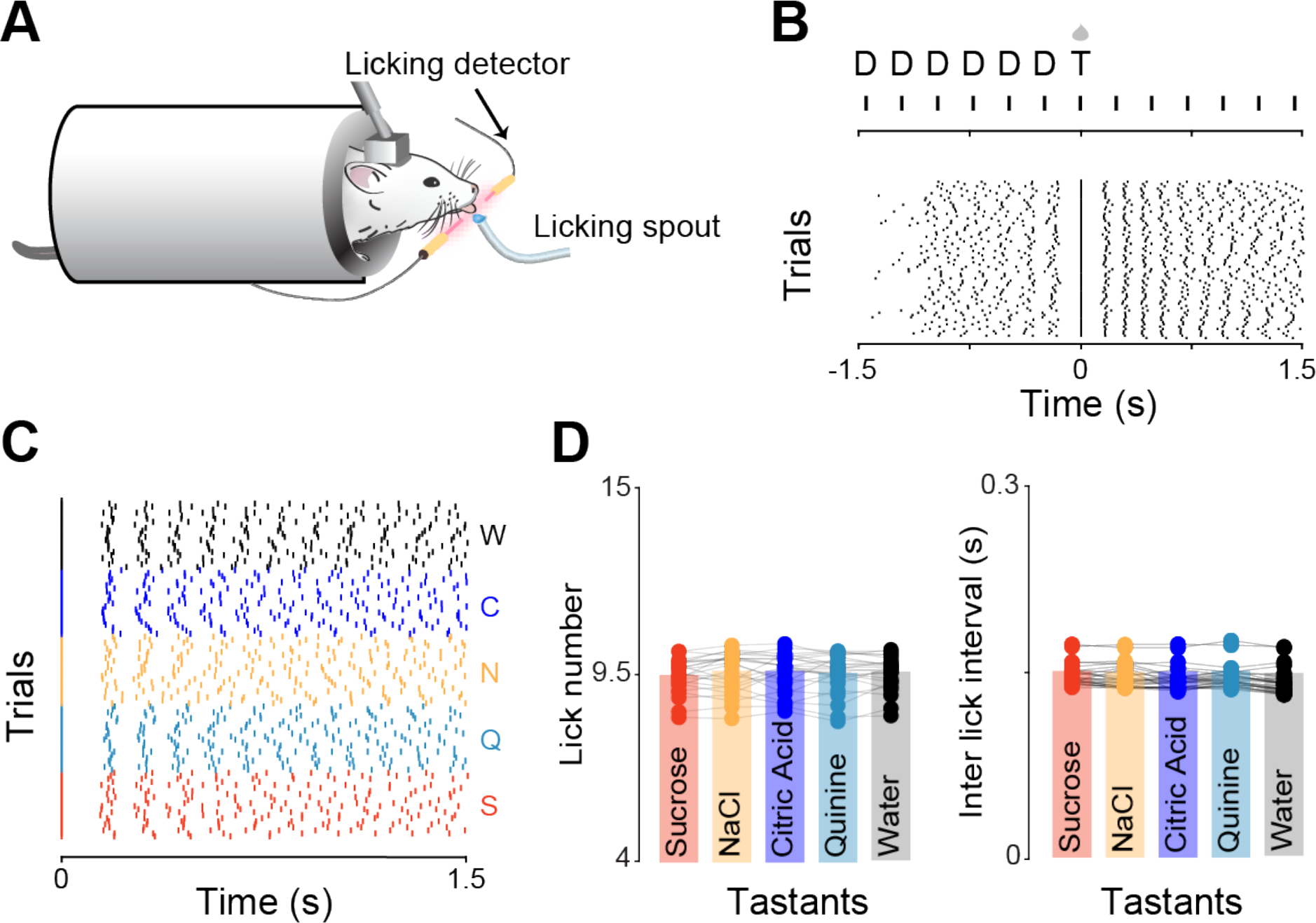
Experimental design and taste-evoked licking pattern. *A*: sketch showing a head-restrained mouse licking a spout to obtain gustatory stimuli. *B*: *top,* diagram of the taste delivery paradigm. Gustatory stimuli (T) are delivered after six consecutive dry licks (D) to the spout. Each lick is symbolized by a vertical line. *Bottom:* raster plot of licking activity in 3 s time interval around taste delivery from one experimental session. Licking activity is realigned to stimulus delivery (time 0 along the x-axis). *C*: raster plot of taste-evoked licking activity. Each line represents an individual lick; trials pertaining to different tastants are grouped together and color-coded with sucrose (S) in red, quinine (Q) in cyan, NaCl (N) in yellow, citric acid (C) in blue, and water (W) in black. *D*: *left,* bar plots showing the number of licks evoked by each gustatory stimulus in a 1.5 s time interval after stimulus; r*ight,* bar plots showing the inter lick intervals (ILI) for the licking pattern evoked by each tastants in a 1.5 s time interval after stimulus. Bars represent the mean, and each round point represents the individual lick number and ILI value extracted for the 25 experimental sessions analyzed.

### Behavioral apparatus and training

Two weeks after full recovery from the surgery (i.e., no sign of distress, proper grooming, proper eating and drinking, return to pre-surgical weight), mice were placed on a water restriction regimen (1.5 ml per day). One week after the start of water restriction, mice were progressively habituated to be head-restrained in the recording rig. The initial duration of head-restraint sessions was short (∼5 min) and gradually increased over days. The restraint apparatus consisted of a metal stage with an elevated clamp for securing the head-bolt. The body of the mouse was covered with a semicircular plastic shelter. The recording sessions took place within a Faraday cage (Type II 36X36X40H CleanBench, TMC), to accommodate for the requirement of electrophysiological recording. Mice were trained with a fixed ratio schedule, in which they learned to lick the dry spout a specific number of times to trigger the delivery of the taste. During training the number of dry spout licks was gradually increased from FR2 (taste stimulus delivered at the 2^nd^ lick) to FR7 (taste stimulus delivered at the 7^th^ lick) before starting the neural recording. The availability of the spout was signaled to the animal by a brief (50 ms) auditory cue. A single trial consisted of the delivery of one out of the five gustatory stimuli followed by a 3*µ*l water rinse 7±1 s after taste delivery. After the delivery of the rinse, an inter-trial interval (ITI) of 6.5±1.5 s separated two consecutive trials. Each recording session consisted of at least of 15 trials per tastant. Water was presented both as a stimulus and as a rinse.

### Electrophysiological recordings

Voltage signals from the tetrodes were acquired, digitized, and band-pass filtered with the Plexon OmniPlex system (Plexon, Dallas, TX) (sampling rate: 40KHz). Single units were offline sorted using a combination of template algorithms, cluster-cutting techniques, and examinations of inter-spike interval plots using Offline Sorter (Plexon, Dallas, TX). Neural data were analyzed with custom-written scripts in Matlab (MathWorks, Inc., Natick, MA). Peristimulus time histograms were plotted around the time of taste delivery. A bin size of 200 ms was used unless otherwise specified.

### Analysis of neural data

#### Stimulus responsiveness

Responsiveness to a taste stimulus or to licking initiation was evaluated by analysis of the firing rate. Concerning the GC neuron sensitivity to the gustatory stimuli, this analysis was first performed by grouping together all of the trials (to provide a general quantification of each neuron’s responsiveness to gustatory stimuli) and then again separating the trials based on the identity of the tastants (to provide further quantification of the responsiveness to each individual taste input). A non-parametric test was used to determine whether the evoked spiking activity (1.5 s after taste delivery to test taste responsiveness and 1 s after the first dry lick to test responsiveness to lick bout) significantly differed from the baseline spiking activity (0.5 s before taste delivery to test taste responsiveness and 0.5 s before the first dry lick to test responsiveness to lick bout). Evoked firing rate was binned in 250 ms-wide temporal windows. Neural activity in each time-bin following the taste stimulus was compared with the spiking activity within 0.5 s of baseline using an imbalanced Wilcoxon rank-sum test with correction for type-I error accomplished by 1) using a Bonferroni correction for multiple comparisons to set the significance cut-off at 0.05/*n* (with *n* = 6) and 2) requiring two-consecutive bins with p < 0.05/*n* to pass. Single units with significant increase of firing rate following the stimulus were deemed excitatory, whereas units with a significant decrease in activity were deemed inhibitory responsive neurons.

#### Taste selectivity

To understand if the spiking activity of GC neurons contained information of specific gustatory stimuli, we quantified their taste-selectivity. Taste-selectivity was assessed by evaluating differences in either the magnitude or time-course of the taste-evoked firing rate across the five stimuli. We employ a two-way ANOVA (taste-identity X time-course) (Samuelsen et al. 2012; Jezzini et al. 2013; Liu and Fontanini 2015; Levitan et al. 2019), using 250 ms bins in the 0 to 1.5 s post-taste time interval. A neuron was deemed taste-selective if the taste-identity main effect or the interaction term (taste-identity X time-course) was significant at p < 0.01.

#### Sharpness index and entropy

To further investigate the response profile of GC neurons, we used sharpness index (SI) (Rainer et al. 1998; Yoshida and Katz 2011) and entropy (H) (Smith and Travers 1979), two standard methods to evaluate taste selectivity used in taste physiology. SI was computed on the mean firing rate during the 1.5 s-wide interval after taste delivery and was defined as:

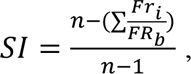

where *Fr_i_* is the mean firing rate for each taste (i = 1–5), *Fr_b_* is the maximum firing rate among gustatory stimuli, and *n* is the total number of stimuli (*n* = 5). A sharpness index of 1 indicated that a neuron responded to one stimulus (narrow tuning), and the value 0 indicated equal responses across stimuli (broad tuning). Entropy metric (H) was computed as previously described by (Smith and Travers 1979):

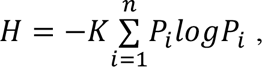

where *K* is a constant (1.43 for 5 taste stimuli) and *P* is the proportional response to each gustatory stimulus (*i*). For this analysis, taste responses were obtained by subtracting the mean taste-evoked firing rate (over 1.5 s after taste delivery) from the mean firing rate preceding taste (over 0.5 s before taste delivery). To control for inhibitory response, the absolute value of taste response was included in the analysis. *P_i_* is the proportional response to each tastant. Overall, a low *H* indicated a narrowly tuned taste-selective neuron, whereas a high *H* indicated a broadly tuned taste-selective neuron.

#### Classification taste identity - population decoding

To understand how well the GC encoded information pertaining the identity of gustatory stimuli and taste information is processed across time, we used a population decoding approach (Meyers 2013). To this end we first constructed a pseudo-population of GC neurons using taste-selective neurons recorded across different sessions (n = 60). We then generated a firing-rate matrix (trials X time-bin) where the spike timestamps of each neuron (1 s before and 1.5 s after taste) were re-aligned to taste delivery, binned into 100 ms time-bins and normalized to Z-score. To assess the amount of taste-related information, we used a “max correlation coefficient” classifier. Spike activity data contained in our matrix were divided into 10 “splits”: 9 (training sets) were used by the classifier algorithm to “learn” the relationship between pattern of neural activity and the different tastants; 1 split (testing set) was used to make predictions about which taste was delivered given the pattern of spiking activity. This process was repeated 10 times (each time using a different training and testing splits) to compute the decoding accuracy, defined as the fraction of trials in which the classifier made correct taste predictions.

#### Palatability-related spiking activity and time-course

To evaluate if GC activity encoded palatability-related information, we used the palatability index (Jezzini et al. 2013). It is important to note that in order to compute the PI, we 1) exclusively considered GC neurons deemed taste-selective by previous analysis (see *Taste selectivity*) and 2) considered spiking activity evoked by sucrose, NaCl, citric acid, and quinine (we did not include water responses in this analysis). To build the PI we considered the time course of the difference of the PSTHs (250 ms bin) in response to tastants of similar (sucrose-NaCl and citric acid-quinine) vs. opposite (sucrose-quinine, sucrose-citric acid, quinine-NaCl, citric acid-NaCl) palatability. To avoid potential confounds introduced by differences in baseline and evoked firing rates across our pools of taste-selective neurons, we first normalized the PSTHs using a receiver operating characteristic (ROC) procedure (Cohen et al. 2012). With this method, the taste-evoked (from time 0 to 1.5 s with time 0 representing taste delivery) firing rate in each time-bin was normalized to the baseline spiking activity (from time -0.5 to 0 s). The normalized firing rate resulted in numbers ranging between zero and one; values larger than 0.5 indicated that the firing rate in that bin is above baseline whereas values below 0.5 indicated that the firing rate is below baseline. After normalizing the firing rate, we then computed the PI. We first computed the absolute value of the log-likelihood ratio of the normalized firing rate for taste responses with similar (<| LR_same_|>) and opposite (<|LR_opposite_|>) hedonic values:

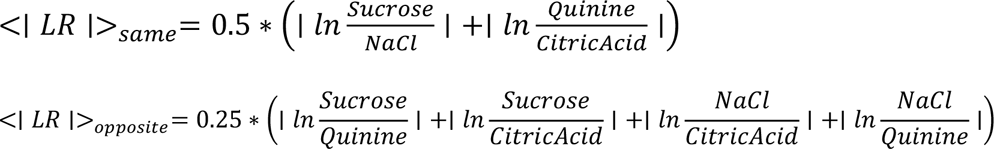

Then we defined the PI as: <|*LR*|>_*opposite*_ − <|*LR*|>_*same*_. Positive PI values (in red in Fig. 8), suggested that a neuron responded similarly to tastants with similar palatability and differently to stimuli with opposite hedonic values. Negative PI (in blue in Fig. 8) values indicated the alternative scenario in which a neuron responded differently to stimuli of same palatability and similar to taste with different hedonic values. A taste-selective GC neuron was deemed palatable-related if its PI value after taste delivery was 1) positive and 2) exceeded the mean + 6*standard deviation of the PI values in the baseline.

#### Licking phase coherence

The coherence (C) between licking and neural (spikes) activity was computed using the “*coherencyc*” function of the Chronux 2.12 software package (http://chronux.org/) (Mitra and Bokil 2008). Multitaper coherence was calculated 1) using licking and neural activity between -1 and 1.5 s (with 0 s being taste delivery; 1 ms bin); 2) using tapers 1-3 and 3) for a frequency range of 5-8 Hz (frequency band observed in freely licking behavior (Spector et al. 1998)). To compute the confidence interval of the coherence and the significant threshold (α at 0.01%) we used a jackknife method (Jarvis and Mitra 2001; Gutierrez et al. 2010); a GC neuron was deemed licking-coherent if its lower confidence interval (99%) crossed the significance threshold.

### Histology

At the end of the experiment, mice were terminally anesthetized and perfused transcardially with 30 ml of PBS followed by 30 ml of 4% paraformaldehyde (PFA). The brains were extracted and post-fixed with PFA for 24 hours, after which coronal brain slices (100 *µ*m thick) containing the GC were sectioned with a vibratome (VT1000 S, Leica). To visualize the tetrodes’ tracks, brain slices were counterstained with Hoechst 33342 (1:5000 dilution, H3570, ThermoFisher, Waltham, MA) using standard techniques and mounted on glass slides. GC sections were viewed and acquired on a confocal microscope (Eclipse Ti2, Nikon).

## RESULTS

To investigate how cortical neurons encode taste information in freely licking mice, we recorded an ensemble of single units via movable bundles of tetrodes implanted unilaterally in the GC (Fig. 1). Tetrode wires were implanted above the GC (AP: 1.2 mm; ML: 3.5 mm; DV: ∼1.1/1.2 mm). Before the first recording session, tetrode bundles were lowered ∼350 *µ*m; after each recording session, tetrodes were further lowered ∼80*µ*m in order to sample new GC neuron ensembles. A total of 283 neurons were isolated across 25 sessions in 6 mice (the average yield was 47 neurons per mouse and 10.4 neurons per session).

One potential challenge of electrophysiological recordings during active sensing is that mice have intrinsic preferences for different tastants. These preferences could result in unique licking patterns for each stimulus – a potential confound for the interpretation of electrophysiological differences. To address this issue, we trained the mice to produce comparable numbers of licks to each stimulus. After habituation to head-restraint, water deprived mice were engaged in a FR7 task in which they had to lick 6 times to a dry spout to obtain a 3*µ*l-drop of one out of five gustatory stimuli (sucrose, 200 mM; NaCl, 50 mM; citric acid, 10 mM; quinine, 0.5 mM; water) (Fig. 2). Mice were trained until the licking pattern evoked by each of the individual gustatory stimuli was similar across a 1.5 s temporal epoch following taste delivery (Fig. 2C). Indeed, no statistical differences in the distribution of total lick number (Fig. 2D, left panel; one way anova, F(4) = 0.13; p = 0.97) and inter-lick interval (Fig. 2D, right panel; one way anova, F(4) = 1.49; p = 0.21) were observed across tastants. The similarity in taste-evoked licking ensured that differences in neural responses across gustatory stimuli within 1.5 s after taste delivery could not be attributed to sensorimotor sources.

### Taste-evoked responses in active licking mice

To begin evaluating the neural dynamics evoked by gustatory stimuli in active licking mice, we analyzed the spiking profile of single GC neurons. Figure 3 shows the raster plots and peristimulus time histograms (PSTHs) of three representative GC neurons. Visual inspection of the graphs indicates that each of these neurons is modulated by at least two gustatory stimuli in a temporally-rich and dynamic manner. In order to understand how many GC neurons were modulated by taste delivery, we compared the firing rate during baseline (averaged between - 0.5 to 0 s) with the evoked spiking activity (averaged between 0 to 1.5 s). Wilcoxon rank sum analysis revealed that a substantial number of the recorded GC neurons (67% [190/283]; referred hereafter as taste-responsive neurons) responded to at least one tastant (Fig. 4A). Specifically, we observed 56% (108/190) of taste-responsive neurons displaying an excitatory response (taste-evoked firing rate > baseline firing rate), while 43% (82/190) displayed an inhibitory response (taste-evoked firing rate < baseline firing rate) (Fig. 4A). Figure 4B shows the population average (population PSTH) of the excitatory and inhibitory response. Analysis of the distribution of the latency of the responses indicated that the majority of taste-responsive neurons showed a fast onset, with firing rate significantly changing from baseline within the first 300 ms after taste delivery (Fig. 4C; mean onset 0.32 ± 0.022 s). The analysis on the duration of the firing rate modulation revealed that the durations of modulation were heterogeneously distributed over the entire 1.5 s post-taste time interval analyzed (mean duration 0.63 ± 0.033 s). Interestingly, no differences pertaining the onset and the duration of the responses were detected between excitatory and inhibitory stimulus-evoked firing rate (onset and duration: Kolmorgov-Smirnov tests, p = 0.8 and p = 0.9, respectively).

**Figure 3.**
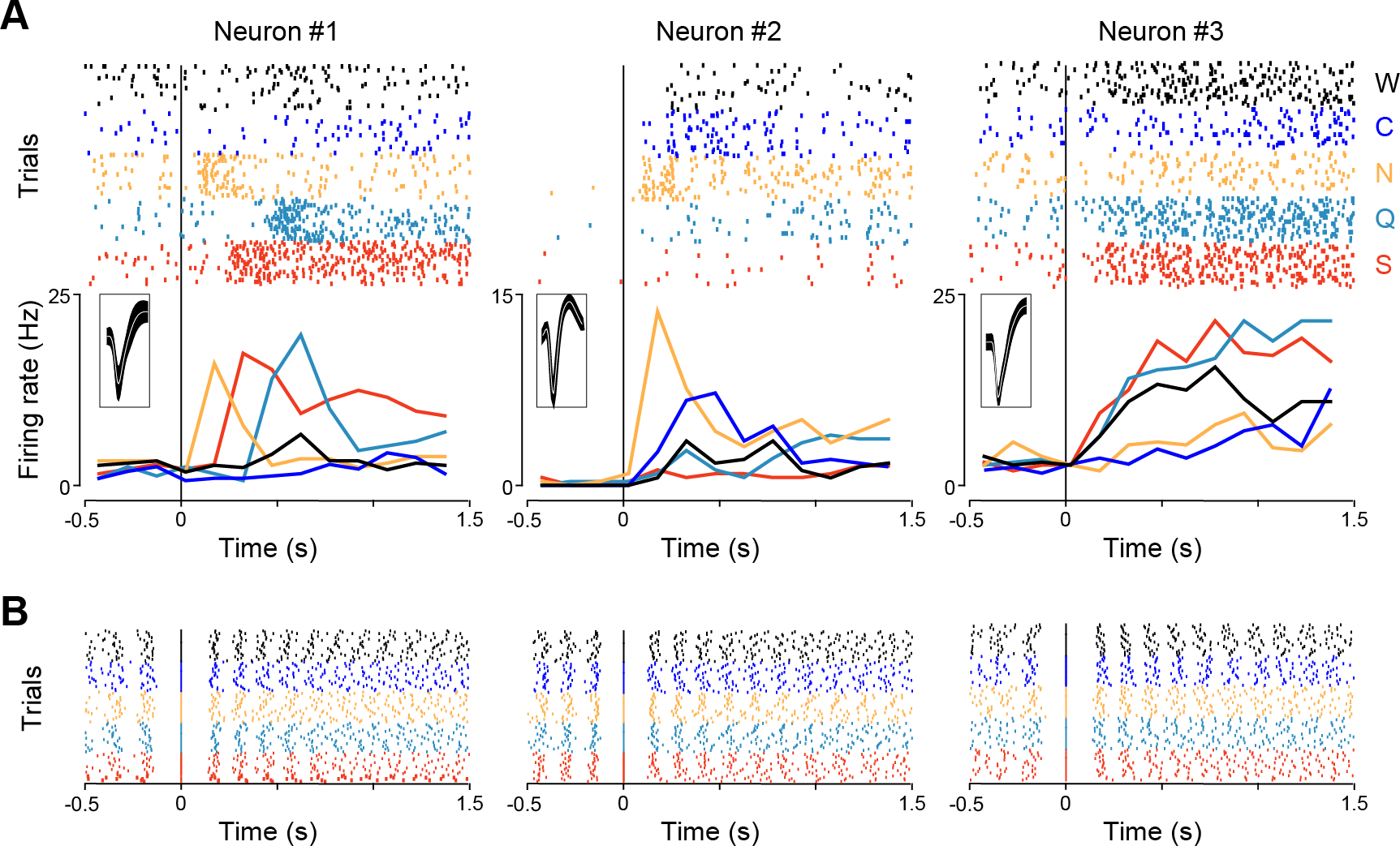
Taste responses in active licking mice. *A:* raster plots and PSTHs of three representative GC neurons showing broadly tuned responses to the five stimuli. Vertical lines at time 0 represent taste delivery. Trials pertaining to different tastants are grouped together (in the raster plots) and color-coded (both in the raster plots and PSTHs) with sucrose (S) in red, quinine (Q) in cyan, NaCl (N) in yellow, citric acid (C) in blue, and water (W) in black. Inset, average action potential waveforms for each of the three neurons. *B:* raster plots of taste-evoked licking activity from the experimental sessions where the neurons shown in A were recorded. Each line represents an individual lick; similar to panel A, trials pertaining to different tastants are grouped together and color-coded with sucrose.

**Figure 4.**
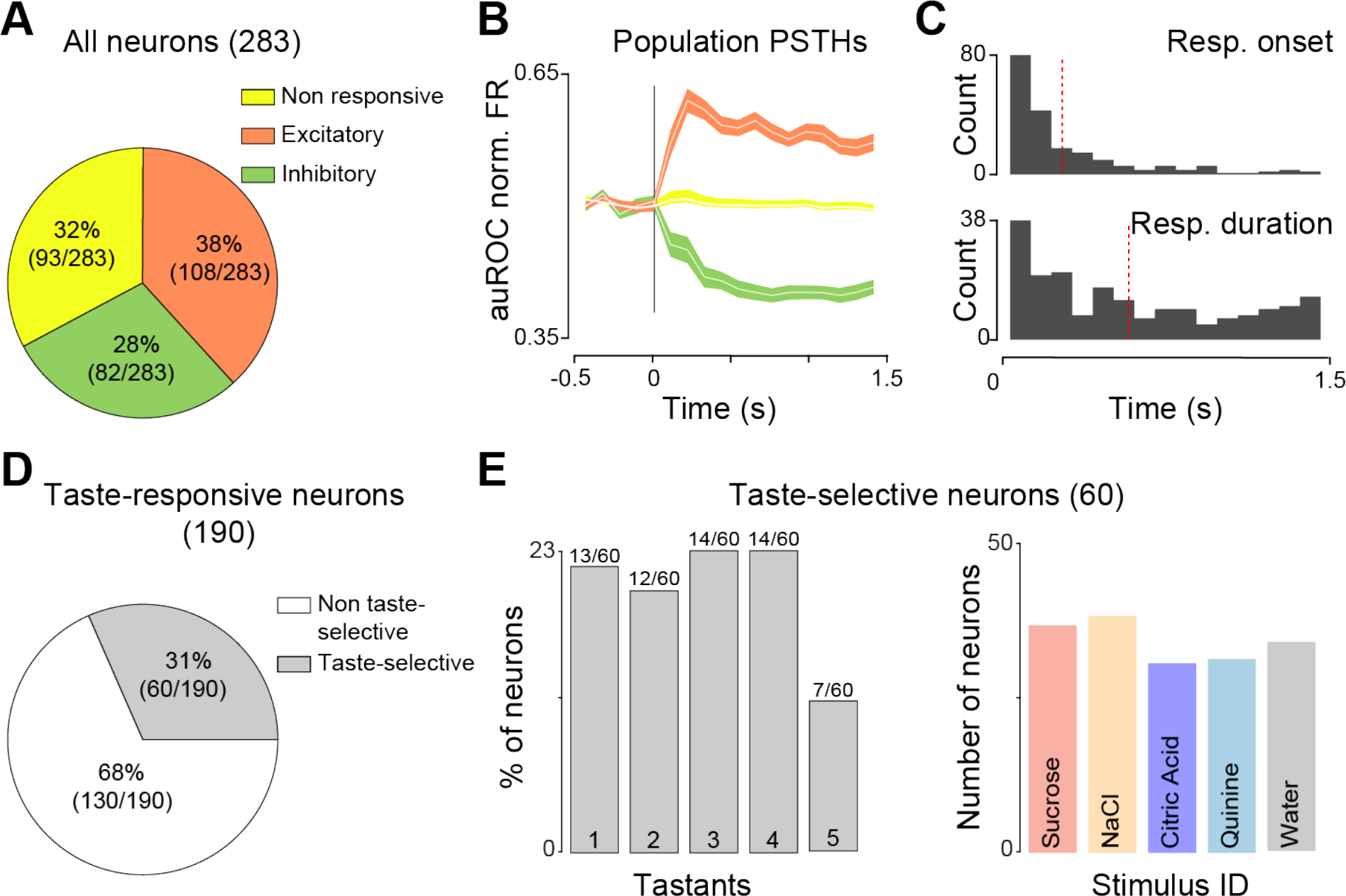
Quantification of taste responses in the mouse GC. *A:* pie chart displaying the proportion and distribution of neurons showing either an excitatory or inhibitory response to taste stimuli. *B:* population PSTHs of excitatory (orange), inhibitory (green) and non-responsive (yellow). Shaded areas represent SEM. *C:* distribution of taste-evoked response onset (top) and response duration (bottom). Vertical red dotted lines represent mean values. D: pie chart showing the proportion of taste-responsive neurons that respond selectively to one or more tastant. Taste selectivity was assessed with two-way Anova comparing responses to the different gustatory stimuli as a function of time. *E*: fraction of taste-selective neurons encoding one, two, three, four, or five gustatory stimuli (on the left) and number of taste-selective neurons responding to the four different taste quality and water (on the right).

### Taste-selectivity and chemosensory tuning of GC neurons

To further understand how the GC processes gustatory information pertaining to the different taste qualities, we moved beyond the taste-responsiveness analysis. In fact, a taste-responsive neuron could be modulated similarly by all stimuli (see for example middle panel in Fig. 6C), reflecting general somatosensory (i.e. the delivery of liquid in the mouth) or cognitive (expectation of a taste based on the FR7 protocol) elements rather than chemosensory specific activity. We address this issue using the following analysis. For each taste-responsive neuron, we compared the spiking activity evoked by the five different gustatory stimuli. Briefly, post-stimulus (0 - 1.5 s) firing rate was divided into six 250-ms bins, and a two-way ANOVA was used with “taste stimuli” and “time” as variables (Jezzini et al. 2013; Levitan et al. 2019). Taste-responsive neurons that had significantly different responses to the 5 tastants in either the main effect “taste stimuli” or the “taste stimuli” by “time” interaction were defined as taste-selective. This method revealed that 31% (60/190) of taste-responsive neurons selectively encoded gustatory information while the remaining (69%) were likely encoding to somatosensory or cognitive features.

**Figure 5.**
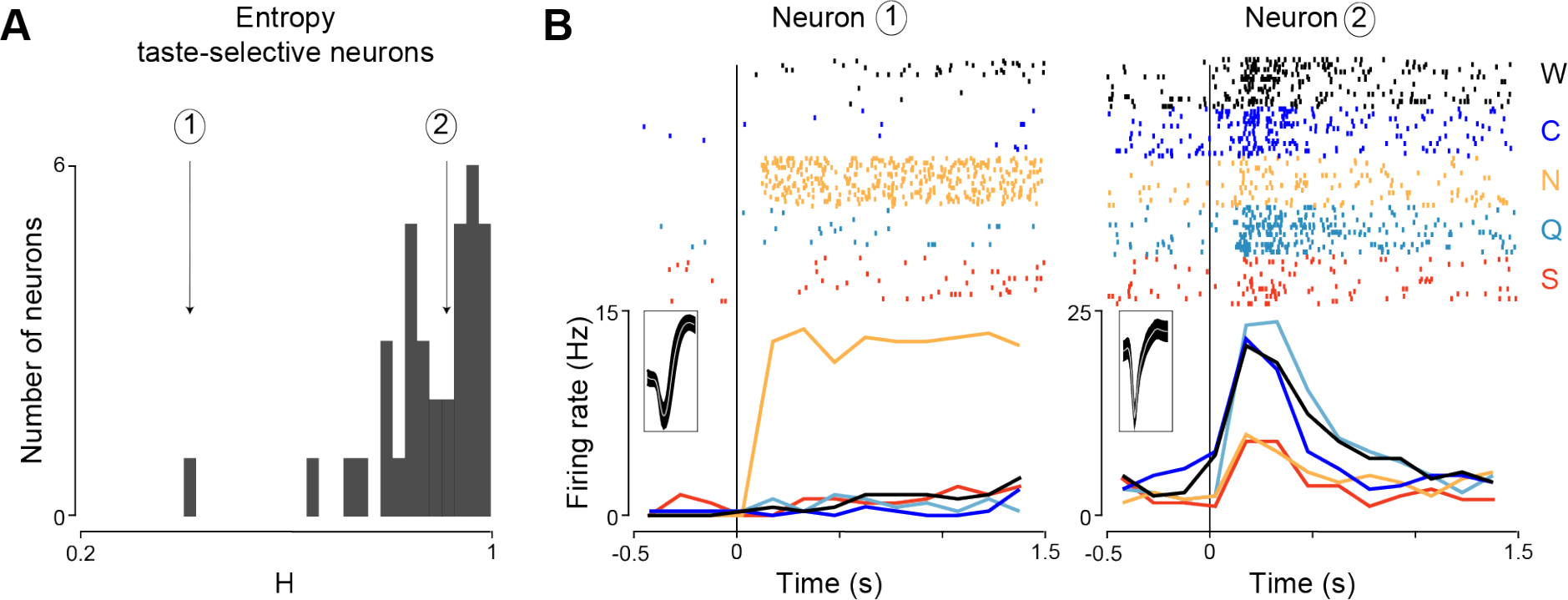
Tuning of taste-selective neurons in the GC of active licking mice. *A:* distribution of the breadth of tuning (expressed as entropy, H) of the taste-selective neurons. Low H values indicate narrowly responsive neurons, while high H values imply that the same neuron is modulated by multiple tastants. The distribution of H values is skewed to the right (toward high H values), indicating that the majority of taste-specific GC neurons are broadly tuned. Black arrows indicate the H values of the two representative neurons shown in B. *B:* raster plots and PSTHs of one narrowly tuned (Neuron 1) and one broadly tuned (Neuron 2) GC neuron. Vertical lines at time 0 represent taste delivery. Trials pertaining to different tastants are grouped together (in the raster plots) and color-coded (both in the raster plots and PSTHs) with sucrose (S), quinine (Q), NaCl (N), citric acid (C), and water (W) in red, cyan, yellow, blue, and black, respectively. The inset figure shows the average spike waveforms for Neuron 1 and 2.

**Figure 6.**
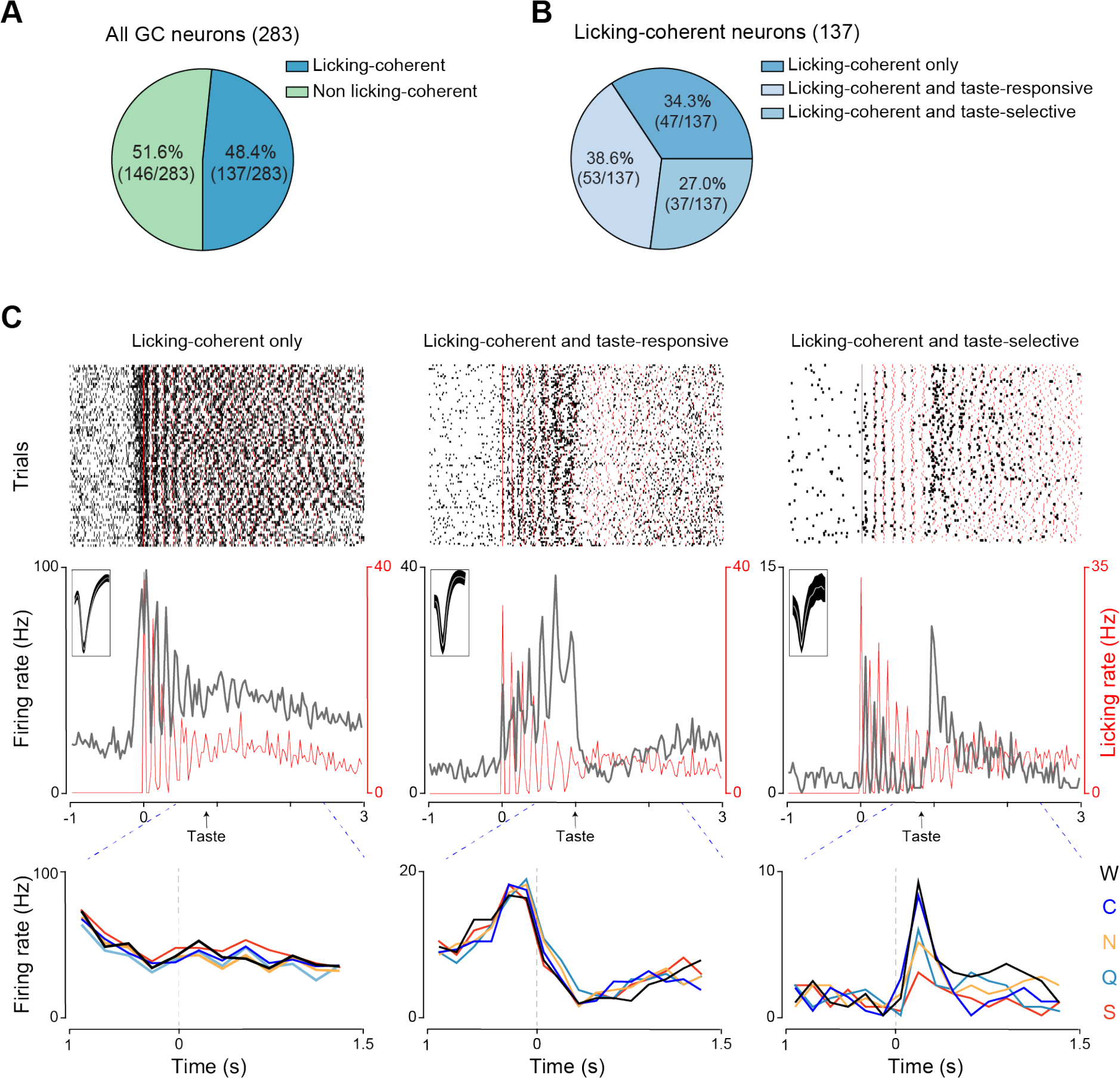
Licking activity in the mouse GC. *A*: pie chart showing the proportion and distribution of GC neurons displaying licking-coherent activity. Note that neurons were defined licking-coherent if they showed significant coherence between licking and neural activity in a temporal window spanning -1 and 1.5 s (with taste delivery at 0 s). *B*: pie chart displaying the proportion of licking-coherent neurons that were also defined as taste-responsive and taste-selective. *C:* raster plots and PSTHs of three representative licking-coherent GC neurons: leftmost panel shows a “licking-coherent only” neuron, central panel a “licking-coherent and taste-responsive” neuron, and the rightmost panel displays a “licking-coherent and taste-specific” neuron. In the raster plots (*top*) each black line represents a spike, and each red line represents a lick; note that in this case trials are not grouped based on the tastants, and neural and licking activities are realigned to the first dry lick. The PSTHs in the central panel represent the average firing (black line) and licking (red line) activity reported in the raster plot, binned in 50 ms time-bins. Neural and licking activities are realigned to the first dry lick (time 0 along the x-axis). The black arrow in the x-axis gives an estimate on when the taste stimulus was delivered. The inset figure shows the average spike waveforms. The PSTHs in the lower panel display the firing rate of the same three neurons. However, spiking activity is realigned to taste delivery (time 0 and gray dotted line in the x-axis), color-coded based on the identity of the gustatory stimulus (sucrose (S) in red, quinine (Q) in cyan, NaCl (N) in yellow, citric acid (C) in blue, and water (W) in black), and binned in 200 ms time-bins.

Next, we evaluated the tuning profile of the taste-selective neurons. We aimed to understand if the taste-selective GC neurons recorded in active licking mice preferentially respond to only one single taste stimulus (i.e. narrow tuning) or if they are capable of encoding information pertaining to multiple tastes (i.e. broad tuning). Our analysis revealed that 21% of GC taste-selective neurons responded to one taste, 20% to two tastes, 23% to three and four tastes, and 11% to all stimuli (Fig. 4B, *left panel*) as shown by the distribution plot in figure 4A. In addition, we observed that taste-selective neurons were not preferentially tuned to encode information pertaining individual taste quality but rather they broadly responded to gustatory stimuli independent of their chemical identity (Fig. 4B, *right panel*). To further investigate differences in the tuning profiles of GC neurons, we performed two additional analyses; we computed the response entropy (*H*) and the response sharpness index (SI) for each taste-selective neuron, two standard techniques used to evaluate the breadth of tuning of single neurons (Smith and Travers 1979; Yoshida and Katz 2011; Wilson and Lemon 2013). Figure 5A shows the distribution of *H* values of the taste-selective neurons. Low *H* values are evidence of narrowly tuned neurons (i.e. GC neurons that encode one taste stimulus; see *Neuron 1* in Fig. 5B) while high *H* values indicate broadly tuned neurons (i.e. GC neurons that encode multiple taste stimuli; see *Neuron 2* in Fig. 5B and the three neurons in Fig. 3). The distribution of H values strongly implies that the majority of taste-selective neurons are broadly tuned, suggesting that the bulk of taste-selective neurons in GC encode information of more than one taste. Similar results were obtained analyzing the response SI (Yoshida and Katz 2011). A sharpness index of 1 describes a neuron responding to only one taste, while a SI of 0 describes a neuron responsive to all 5 gustatory stimuli. The results of this analysis further confirmed that GC neurons are broadly tuned with an average SI of 0.34 ± 0.02 (*data not shown*).

Altogether our analyses revealed that, 30% of GC neurons in active licking mice were capable of selectively encoding taste identity information. In addition, we showed that the majority of these neurons displayed a broad tuned profile, indicating that a single taste-selective neuron is likely to process information of different gustatory stimuli.

### Licking-coherent neurons

Electrophysiological studies performed in rats have indicated that neurons in the GC can integrate somatosensory and motor activity linked to licking (Gutierrez et al. 2010; Stapleton et al. 2006). To determine how rhythmic licking impacted neural activity in the mouse GC, we quantified the number of neurons that exhibited neural activity coherent with licking. In our analysis, coherence (C) revealed how well the spiking activity of one neuron was correlated with rhythmic licking in the 5-8 Hz frequency domain (see material and methods for further details). This analysis revealed that a substantial amount of GC neurons (137/283, 48.4% referred hereafter licking-coherent; Fig. 6A) exhibited spiking activity time-locked and correlated to subsequent licks in both frequency and phase. We examined whether the licking-coherent neurons were exclusively modulated by lick-related activity or if they also encode taste-related chemosensory information. Additional analyses on the firing rate modulation before and after taste delivery revealed three types of licking-coherent neurons (Fig. 6B). 34.3% (47/137) of the total licking-coherent neurons (Fig. 6B, leftmost panel, *licking-coherent only*) displayed a rhythmic and lick-sensitive spiking activity with no differences in the firing rates evoked by licks preceding and following taste delivery and across tastants. Likely, these neurons exclusively encode somatosensory and/or motor aspects of licking and do not integrate the information pertaining tactile inputs from fluids delivery in the oral cavity nor the chemosensory identity of tastants. 38.6% (Fig. 6B, *licking coherent and taste responsive*; 53/137) of licking-coherent neurons displayed a clear change in spiking activity following taste delivery but no differences across tastants. These neurons likely encode both somatosensory-motor features of licking and the presence of a taste solution in the mouth independent on taste quality and identity. The remaining 27% (Fig. 6C, *licking coherent and taste selective* 37/137) of neurons not only displayed lick- and taste-sensitive activity but their taste-evoked firing rate selectively discriminated for the chemosensory identity of the tastants. A test for equality of proportion revealed that the fraction of licking-coherent GC neurons are similar across the three types (“licking-coherent only” 34.3% [47/137], “licking-coherent and taste responsive” 38.6% [53/137] and “licking-coherent and taste-selective” 27.0% [37/137]; proportion test: **χ**^2^_(2)_ = 4.29; p = 0.11).

Together, these results indicate that half of the neurons recorded in the GC of active licking mice are strongly correlated with licking activity. While the majority of these licking-coherent neurons are lick-sensitive and encode somatosensory, yet chemosensory-independent information of a fluid in the mouth, a substantial fraction was found to be taste-selective. Indeed, analysis of the temporal dynamics of the taste-evoked firing rate in 1.5 s after taste delivery, revealed that 27% of licking coherent neurons also encode chemosensory information.

### Population decoding of taste information in active licking mice

After characterizing the profile of the chemosensory and licking-related responses in single neurons, we focused our attention on the neural activity at population level. While single neuron activity can encode important features of sensory stimuli, information encoded in networks (population or ensemble) of neurons is used to inform behavioral choices. For example, sudden changes in taste-evoked activity of GC neuron ensembles correlate with (Stapleton et al. 2006; Gutierrez et al. 2010) and are needed to drive (Mukherjee et al. 2019) behaviors (gapes in this case) aimed at expelling highly unpalatable tastants. Therefore, we used a population decoding analysis to quantify the amount of gustatory information contained in the firing patterns of ensembles of GC neurons.

We computed the taste decoding performances in our taste-selective neurons (i.e. neurons whose firing rate selectively discriminated for the chemosensory identity of the tastants; n = 60; gold trace in Fig. 7A). The time course of the taste classification average across tastants is shown in Fig. 7A. Taste decoding showed an early onset (classification above chance from the first bin after taste delivery) and reached its peak around 500 ms after taste delivery. In addition, while the overall classification value started decreasing after 500 ms, decoding performances remained above chance until the end of the temporal window analyzed (Fig. 7A). As a control, the same analysis was performed using non taste-selective neurons (neurons that responded to all gustatory stimuli similarly; n = 130; black trace in Fig. 7A). As expected, the taste classification accuracy never exceeded chance level (Fig. 7A). The time-course of the classification analysis suggested that the spiking activity of GC neurons contained an optimal amount of information to discriminate the different gustatory stimuli in 500 ms (∼ four licks) after taste delivery. However, it is well established that rodents are able to identify and discriminate the identity of gustatory stimuli in a single lick (∼120 ms) (Halpern and Tapper 1971; Graham et al. 2014). To evaluate taste decoding at a finer time-scale, we constructed confusion matrices and characterized the classification performance for each taste in 120 ms time-bins around taste delivery (Fig. 7B). As Figure 7A shows the average decoding performance in taste-selective neurons is well above chance (20%, black line in color bar in Fig. 7B; dark-yellow color) during the first lick (between 0 and 120 ms). However, inspection of the confusion matrices in Fig. 7B revealed that, in this early phase of taste processing, not all taste stimuli are equally classified (*χ*^2^_(4)_ = 63.36; p < 0.001). In this plot, the main diagonal highlights the fraction of trials in which the classifier correctly assigned the taste stimulus (predicted taste) to its real category (true taste).Comparison of the fraction of trials correctly classified for each individual tastant in the first 120 ms revealed that NaCl is the tastant best predicted by the decoding algorithm (Marascuillo’s test, p < 0.01). As time progressed, all gustatory stimuli were similarly decoded with > 40% accuracy (well above chance which is 20%) within 320 ms (*χ*^2^_(4)_ = 8.68; p = 0.07).

**Figure 7.**
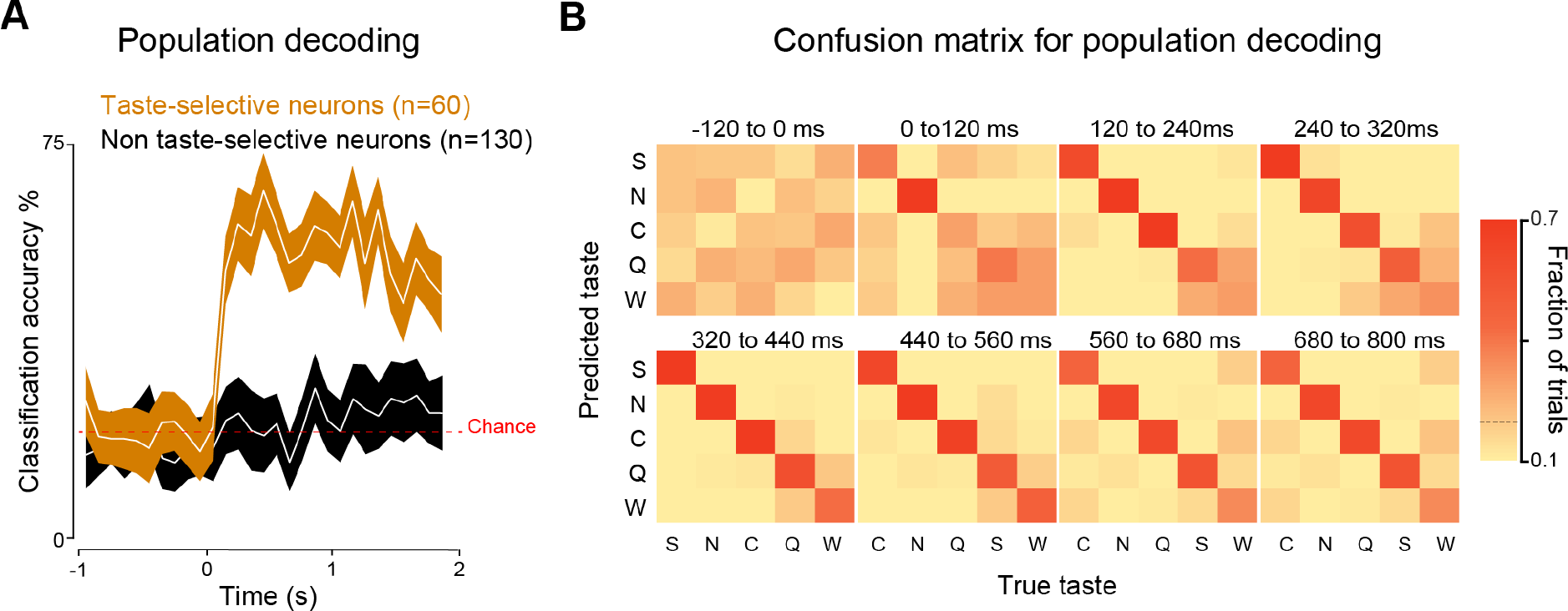
Population decoding analysis and time-course. *A*: time-course of decoding performance considering the population of taste-selective (gold, n = 60) and non-selective (black, n = 130) neurons. Shading represents the bootstrapped CI, while red dotted line indicates chance level performance (20%). Time 0 indicates taste stimulus delivery. Note that the temporal evolution of the spiking activity of the non selective neurons does not convey information about taste. *B*: confusion matrix showing decoding performance for each gustatory stimuli in different 120 ms temporal windows around taste delivery (0 s). Color codes the classification accuracy, with darker hues indicating a higher fraction of correct trials. The main diagonal highlights the number of trials in which the classifier correctly assigned the taste stimulus (predicted taste) to its real category (true taste).

**Figure 8.**
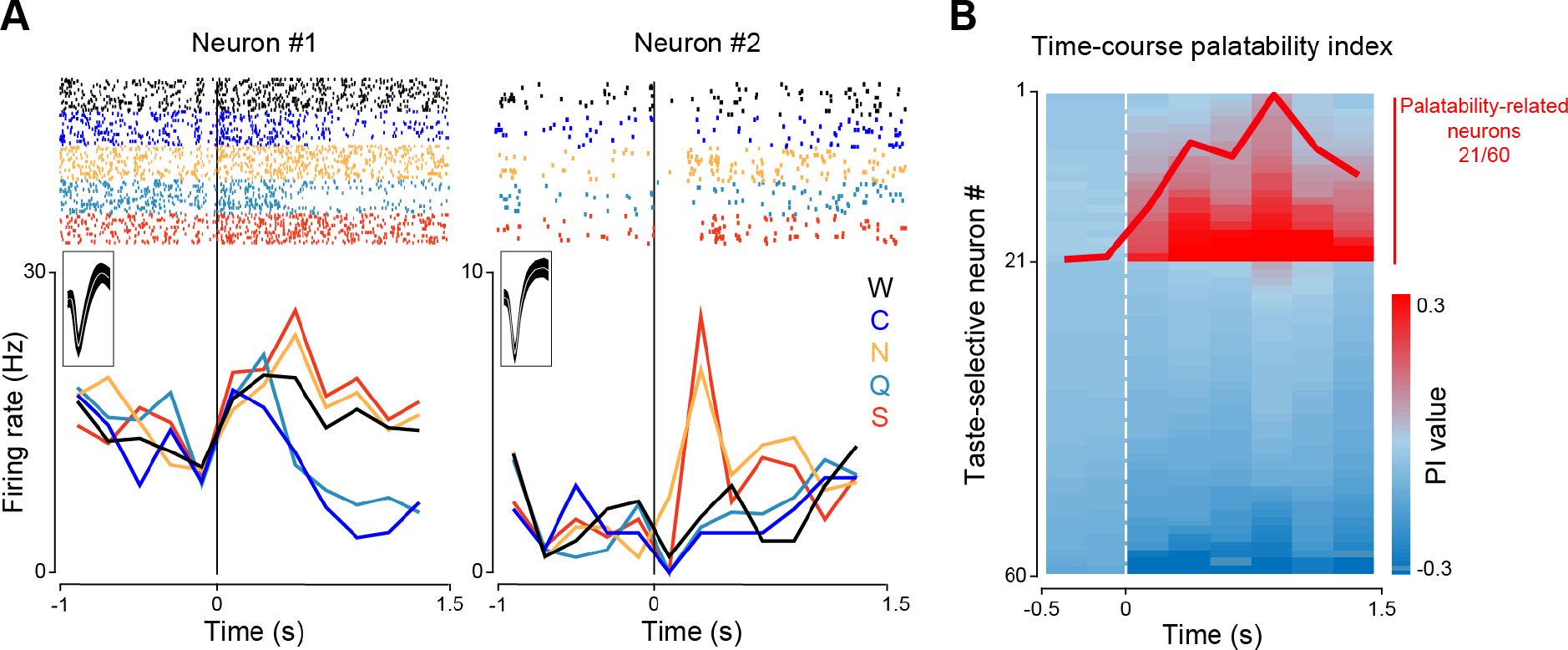
Hedonic processing in the mouse GC. *A*: raster plots and PSTHs of two neurons displaying palatable-related spiking activity. Vertical lines at time 0 represent taste delivery. Trials pertaining to different tastants are grouped together (in the raster plots) and color-coded (both in the raster plots and PSTHs) with sucrose (S) in red, quinine (Q) in cyan, NaCl (N) in yellow, citric acid (C) in blue, and water (W) in black. The inset figure shows the average spike waveforms. *B*: color-coded plot showing the palatability index (PI) values across 2 s around taste delivery at 0 s for all 60 taste-selective neurons. Each row represents a single neuron; time 0 and white vertical dotted lines highlight taste delivery. The thick red line superimposed on the color map represents the time-course of the PI average value for the 21 taste-selective neurons that are deemed palatability-related.

Altogether, these data indicate that ensembles of GC neurons recorded from active licking mice reliably encoded chemosensory information up to 2 seconds after taste delivery. In addition, taste coding showed an early onset, with the time-course of the population decoding performance rising above chance within the first lick (∼120 ms) and peaking after three consecutive licks (∼320 ms).

### Processing of palatability-related taste information in active licking mice

Studies in both anesthetized and alert rodents revealed that GC neurons also encode information about taste palatability when tastants are delivery intraorally (Katz et al. 2001; Accolla and Carleton 2008; Sadacca et al. 2012; Jezzini et al. 2013; Levitan et al. 2019).

However, it is still unknown if GC neurons are capable of encoding hedonic information when gustatory stimuli are actively sensed through licking. Visual inspection of spiking activity in Fig. 8 indicates the presence of palatable-related neurons (see also Neuron 2 in Fig. 5) with both raster plots and PSTHs highlighting the similarity of the firing rate evoked by tastants belonging to the same hedonic category. To quantify the numbers of GC neurons encoding palatability-related features, we computed a palatability index (Jezzini et al. 2013; Liu and Fontanini 2015) (PI; see material and methods for further details). This analysis was based on extracting the taste-response similarity for each taste-selective neurons (n = 60): positive PI values indicate palatability coding, and negative PI values imply inverse palatability coding (similar activity for tastants with opposite hedonic value and different activity for tastants with similar hedonic value). Using this analysis, a neuron was deemed as encoding palatability if 1) it had a positive PI value (it responded similarly to gustatory stimuli of similar hedonic value [sucrose-NaCl and citric acid-quinine)] and differently to tastants with opposite palatability [sucrose-citric acid, sucrose-quinine, NaCl-quinine and NaCl-citric acid]), and 2) it had a PI significantly above baseline for at least 250 ms. This analysis revealed that 35% (21/60) of taste-selective neurons encoded hedonic value (Fig. 8A). In order to determine the temporal evolution of the neural processing of palatability, we analyzed the time course of the PI index (Fig. 8B). The thick red line overlaying the heat map plot shows the PI averaged across palatability coding neurons. Palatability coding showed an early onset with two peaks (highest significant PI values): one before 500 ms and one before 1s after taste delivery. As a control, the PI time-course of taste-selective neurons not coding for hedonic values is also plotted and showed no modulation.

Overall, these results demonstrate that neurons in the mouse GC can encode taste palatability in addition to chemosensory identity. Interestingly, in active licking mice, GC hedonic coding emerges rapidly in the first second after taste delivery.

## DISCUSSION

The results presented here provide evidence on how the gustatory cortex (GC) of mice encodes taste information when stimuli are sampled via active licking. Tetrode recordings revealed that GC neurons process information pertaining to the chemosensory identity of tastants in a temporally dynamic manner. Around 30% of the recorded neurons displayed taste-specific spiking activity, with the majority encoding for more than one taste. GC neurons also processed information of the hedonic value of tastants. Stimuli with similar palatability (i.e. sucrose and NaCl; citric acid and quinine) evoked similar firing activity in more than half of the taste-selective neurons. Analysis of the temporal sequence of both taste identity and hedonic value revealed that chemosensory information is encoded first, followed by palatability. Interestingly, coding of both features of gustatory stimuli occurred rapidly. Decoding population analysis revealed that taste identity started to be encoded during the first lick (within 120 ms after taste delivery), and palatability information arose and peaked within ∼800 ms after taste delivery. Our data also revealed that the neural activity of nearly half of the GC neurons was coherent with licking, likely processing general tactile and motor activity. Notably, almost 30% of licking-coherent neurons were also taste-specific, highlighting the capability of the GC to multiplex different features of taste experience.

### Chemosensory and palatability neurons in the GC of active licking mice

To date, only a handful of studies have investigated the electrophysiological profile and taste-evoked activity of the taste cortex in alert mice. Single neuron spiking activity in response to gustatory stimuli has been examined in awake mice, either using a limited set of tastants and focusing on brief post-stimulus interval (Kusumoto-Yoshida et al. 2015; Vincis et al. 2019) or by flushing taste stimuli directly in the oral cavity via surgically implanted intraoral cannulae (IOC) (Levitan et al. 2019). This study represents the first electrophysiological investigation of the taste-evoked activity over a long post-stimulus interval of GC neurons recorded from actively licking mice. Our experiments show that 66% of the recorded neurons were modulated by at least one taste with a substantial fraction (31% of them; referred as taste-selective neurons) capable of selectively encoding taste identity. Analysis of the breadth of tuning revealed that the majority of taste-selective neurons were broadly tuned, responding to more than one tastant. Most significantly, our results were in agreement with the conclusions reached in recent studies in which gustatory stimuli were delivered directly into the oral cavity. In fact, both *in-vivo* calcium imaging (Fletcher et al. 2017; Livneh et al. 2017) and electrophysiological recordings from awake mice (Levitan et al. 2019) have indicated that a substantial fraction of GC neurons are modulated by gustatory stimuli and are likely to respond in a broadly tuned manner.

Previous electrophysiological data have shown that GC neurons not only encode the physiochemical properties of tastants (i.e. their chemosensory identity) but also code for palatability (Katz et al. 2001; Mukherjee et al. 2019; Levitan et al. 2019). However, these studies used tastants delivered via IOC. We evaluated whether GC neurons encode tastant hedonic value even when they are actively sampled via licking.

It is important to note that as a consequence of our experimental design, there are at least two caveats that can potentially mask palatability-related neural activity. First, compared to other studies (see Levitan et al. 2019), we used taste concentrations that were two times more diluted (at least for three out of four tastants), thus reducing the range of palatability. Second, we recorded neural activity only from mice that displayed a taste-evoked licking microstructure (often use to behaviorally assess taste palatability) that was similar across stimuli after 7-10 days of training (see Material and Method). Nevertheless, relying on a firing rate analysis method used extensively in the field [palatability index, (Piette et al. 2012; Jezzini et al. 2013; Liu and Fontanini 2015)], we determined that 35% of taste-selective neurons process information pertaining the hedonic value of the taste. While our experiments were not directly designed to investigate variations in taste coding between rodent species (rats *vs.* mice) or taste delivery methods (IOCs *vs.* active licking), it is interesting to discuss similarities and potential differences. Our results suggest that the GC processes tastants similarly in both mice (this study) and rats, regardless of whether the tastants were actively sampled (this study) or delivered via IOC (Katz et al. 2001; Jezzini et al. 2013; Levitan et al. 2019). Overall, GC neurons appear to use a common encoding strategy to process identity, including broadly tuned neurons, and palatability features of tastants regardless of rodent species and taste-delivery methods (see *temporal processing* and *lick-related activity*, below).

### Temporal processing of taste coding in active licking mice

Cortical taste processing is characterized by a dynamic and time-varying modulation of the firing activity of GC neurons (Katz et al. 2001). Various studies (in rats and mice) have extensively described and validated a model illustrating that taste responses evolve during multiple distinct temporal epochs in a 2 s time-span following taste delivery (Katz et al. 2001; Grossman et al. 2008; Levitan et al. 2019). The first 200 ms were described as processing “general” tactile information of fluids contacting the oral cavity and were followed by two other temporal epochs in which chemosensory (200 ms - 1 s after stimulus delivery) and palatability (> 1 s after stimulus delivery) features of taste were sequentially encoded. An outstanding question in the field is whether a similar temporal evolution of taste processing is observed during active sensing. Here we focused on the temporal dynamics of chemosensory coding. Population decoding revealed that GC neurons encode taste information in a single lick (Fig. 7). Our analysis indicated that the spiking activity of GC neurons contained sufficient information to discriminate among tastants in the first 120 ms after stimulus delivery.

Noteworthy to mention, while all tastants were classified with great accuracy (>50%) within the first 320 ms, our data suggested that the GC encodes NaCl with the shortest latency (Fig. 7B). Our data seems to indicate that GC neurons encode chemosensory information more rapidly when tastants are actively sampled through licking compared to when they are flushed passively into the mouth via IOCs (Katz et al. 2001). What could explain this difference? Tastants delivered via IOCs are often delivered to the mouth of the animal unexpectedly. In our experimental setting, the licking spout and the fixed ratio paradigm could serve as anticipatory cues, allowing the animal to predict (i.e. expect) that a tastant would be delivered. Interestingly, a previous study has also shown that GC coding of gustatory stimuli can be expedited with anticipatory cues, priming the cortex for faster processing of chemosensory information (Samuelsen et al. 2012). This leads us to reason that actively sampled tastants are intrinsically expected, therefore prompting coding within the GC to occur faster than when tastants are unexpectedly delivered in the animal’s mouth.

We then directed our attention to the time-course of palatability coding. Further analysis of the temporal dynamics of taste-evoked firing rates reported overall rapid and persistent palatability processing. The time-course of hedonic coding revealed two distinguishable palatability phases: a first “early” epoch that peaked at ∼500 ms and a second “late” epoch peaking ∼1 s after taste delivery. It is important to note that some of the GC neurons encoding palatability displayed both phases (like neuron #2 in figure 8), suggesting that the GC receives two “waves” of hedonic information that can converge on the same neuron. While the early phase of hedonic coding reported in our study precedes the appearance of palatability information obtained from neural recordings in the GC of rats (Katz et al. 2001; Jezzini et al. 2013), it does corroborate recent observations in the GC of mice receiving tastants passively from IOCs (Levitan et al. 2019). Overall, our experiments reaffirm and expand our knowledge on cortical taste processing, demonstrating that temporal multiplexing of gustatory information can be observed also in the context of mice sampling tastants via active licking.

### Lick-related neural activity in the mouse GC

Taste perception arises from the association between gustatory and oral-somatosensory information originating from receptors located in the oral cavity (Simon et al. 2006). The anatomical proximity and the neural connections between the gustatory and somatosensory cortices suggest that taste and oral somesthetic information could be intermingled in the cerebral cortex. Indeed, multiple studies in rats (Katz et al. 2001; Stapleton et al. 2006; Gutierrez et al. 2010) have shown that GC neurons displaying a characteristic firing rate in the 5-10 Hz frequency domain can process lick-related somatosensory activity. These results suggest that the majority of licking-coherent neurons fail to discriminate between the different tastants, suggesting that oral chemosensory and somatosensory information are processed by different groups of GC neurons. Our results confirm and expand this view. Analysis of the neural responses to licking activity and gustatory stimuli revealed that the majority of licking-coherent GC neurons failed to discriminate chemosensory information. As was already shown in subcortical taste-related brain areas (Roussin et al. 2012; Weiss et al. 2014), these neurons are likely conveying either rhythmic licking information or general intraoral tactile features and might collaborate with chemosensory neurons to identify and process taste information. However, our data indicate that not all of the GC licking-coherent neurons belong to the aforementioned category. Indeed, almost one-third of the neurons displaying spiking activity that correlated with licks also encoded taste information (Fig. 6). These data support the possibility that GC neurons received multimodal input from both chemo- and mechanosensory projections in the oral cavity.

## ACKNOWLEDGMENTS

The authors would like to acknowledge Dr. Douglas Storace, Dr. Alfredo Fontanini and the members of the Vincis laboratory for their feedback and insightful comments. This work has been supported by National Institute on Deafness and Other Communication Disorders Grant R21-DC016714.

## AUTHOR CONTRIBUTION

C.B. and R.V. carried out study conceptualization and experimental design. C.B. and R.V. performed experiments and analysis. C.B. and R.V. wrote the manuscript.

## DECLARATION OF INTERESTS

The authors declare no competing interests.

